# Comprehensive BioImaging Study of the Red Permanent Marker Ink: Re-purposing for Cells Imaging Including Cytoplasmic Membrane Visualization and Comparison with Rhodamine 6G, Deep Red Cell Mask, and DiBAC^⋆^

**DOI:** 10.64898/2026.04.13.717455

**Authors:** Anna A. Abelit, Natalia A. Boitsova, Liudmila E. Yakovleva, Anton A. Kornev, Daniil D. Stupin

## Abstract

In this paper, we aim to present a new intravital cells visualization method, which is based on use of a dye called ABDS (“A Beautiful dye for staining”), which can be prepared using a marker pen and is useful for eukaryotic cell research. Using a wide range of instruments, including optical measurements, microscopy studies and wet biology techniques, we have shown that ABDS is close by properties to Rhodamine 6G dye (R6G), which is well known as endoplasmic reticulum stainer. However, by the careful examination of the ABDS and R6G images (ABDS/R6G), we have proved for the first time that these dyes also stain the cytoplasmic membranes. The significant contrast between ABDS/R6G signal from cell membrane and endoplasmic reticulum allows them to be distinguished in the fluorescence photographs. Other important properties of ABDS are its availability, simplicity in manufacturing, safety for living cells *in vitro*, and bright stable fluorescence, which in contrast to commercial dye like DiBAC allows us to study cells in space and time with high detalization. The paper includes a method for preparing ABDS, a data set with its characteristics, comparison with other commercial dyes, as well as examples of ABDS usage in cells research.

**Graphical Abstract:** 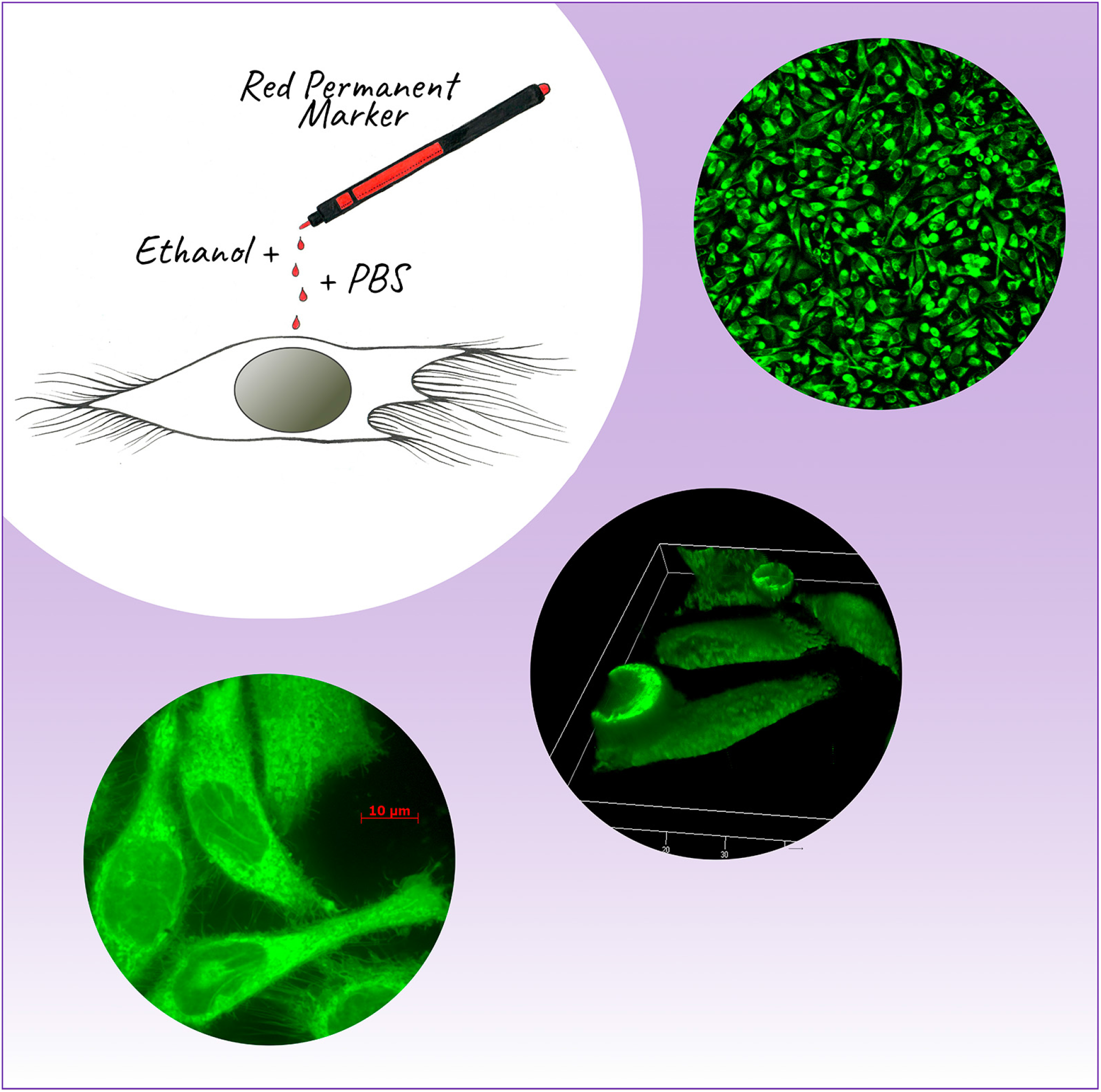

**Highlights:**

- A protocol for high-resolution vital staining of the cells using an inexpensive dye based on permanent marker ink is proposed.
- The absorption, emission and Raman spectra of the proposed dye are presented, and a direct comparison with commercial dyes Rhodamine 6G, DiBAC and Deep Red Cell Mask dye is made.
- The main characteristics of the proposed dye are low toxicity, long-term fluorescence, and the ability to separately stain the endoplasmic reticulum and cytoplasmic membrane.
- The ability of the Rhodamine 6G dye to stain cell membranes also has been proved.

## 1. Introduction

The history of science shows that important discoveries in the modern era of high technologies can still be made using mundane things and materials, which are applied in an unexpected way. Famous examples include Fleming’s invention of penicillin with an accidental poorly washed dishes [1]; the discovery of artificial radioactivity supported by Fermi’s neutron-slowing aquarium [2]; the invention of X-ray film medical technique by Röntgen, who observed the bones of his wife’s hands on a photographic plate [3]; and finally, graphene, discovered with scotch tape and a graphite pencil in 2004 by Geim and Novoselov [4]. As a matter of fact, in ahead in development fields of human activity the ability to use improvised means also should not be underestimated, as was brightly shown in the famous “Houston, we have a problem”-accident on board the Apollo 13 spacecraft in 1970, when the engineers on Earth had to come up with a way to fix the malfunction using items on hand at the astronauts’ disposal [5].

Even in the advantage research of modern biology, there is a place for using everyday things. For example, nail polish is useful for fixing coverslips for microscopic and fluorescence samples [6, 7, 8, 9], a gas torch can be used to sterilize metal surfaces [10], and non-fat milk is used in biochemistry protocols [11, 12]. Another striking example is the use of mobile games – an inseparable part of everyday life – to assist researches related to cancer (Cancer Research UK games, Genima) [13, 14] and protein folding (FoldIt) [15]. The latter program is notable for inspiring researchers to use artificial intelligence to predict protein structure, an approach that was awarded a Nobel Prize in 2024 [16].

Frequently, scientists’ motivation to improvise with methods and materials at hand is caused by a desire to increase research cost-efficiency, speed up study progress, or due to the absence of available on the market required chemicals or apparatus [17]. Moreover, as noted in the remarkable work [18], the use of do-it-yourself (DIY) and lab-in-box technologies allows scientific activity to be carried out even in developing countries, as well as in areas of epidemics and difficult-to-access regions.

One of the bright examples of DIY research optimization in biology was done in 2013 by Yuta Takase *et al*. who have used highlighter ink to visualize blood vessels in avian and mouse embryos using fluorescent microscopy [19]. In 2021, the method of visualizing blood vessels using yellow highlighter ink was used in a study by Luke Bollinger and Renee Dickie, who studied axolotl regeneration [20]. They also improved this imaging method by making it possible to image vessels using a fluorescent adapter system with a stereomicroscope and a phone camera. Finally, visualizing vessels using highlighter ink was also used in a 2023 by Mijithra Ganesan *et al*., in their study of regeneration in earthworms [21]. This study did not use fluorescence microscopy, but visualized vessels using highlighter ink, allowing them to be observed with a light microscope.

In our study, following this line, we expand the area of such a “lab-scale” concept by improving the idea of the writing implements utilization in biology, and propose a cell membrane and endoplasmic reticulum vital dye, which is prepared from commercially available permanent marker ink. Due to the outstanding property of cells visualization, we named our dye ABDS as the abbreviation of “A Beautiful dye for staining”. Contrary to works [19, 20, 21] our staining technique allows to investigate micro-objects such as single living cells in real time mode with submicron resolution. The search for new types of cell dyes and cell staining techniques is very important for fluorescence microscopy, which is used to study processes that occur in cells and for drug testing [22, 23, 24, 25].

We present (i) the preparation protocol for ABDS including a 3D printed stencil to control the concentration of ABDS, (ii) the study of the optical properties of ABDS including absorption, fluorescence and Raman spectra and photobleaching dynamics, (iii) investigation of the ABDS labeling regions and ABDS cytotoxicity and phototoxicity, (iv) the demonstration of the applications of ABDS in living cell research, including the comparison of ABDS with a low-cost cell membrane dye DiBAC (ThermoFisher Scientific, USA). Using the obtained data, we have drawn conclusions about ABDS composition and found the effect of gradual increase in brightness of ABDS during long-term optical pumping, which can be explained by quantum mechanics. Furthermore, in addition to the fluorescence rising effect, ABDS exhibits significantly longer fluorescence intensity conservation with respect to DiBAC dye, allowing for clear, high-quality z-stack and time-lapse images of cells without typical bleaching artifacts.

We also showed that ABDS images of cells contain, among other things, a signal from the *cytoplasmic membrane*, and proposed a method for visualizing the cell membrane using ABDS staining and subsequent software processing. We had also for the first time observed the similar cell membrane staining effect of the Rhodamine 6G, the highly possible staining component of the ABDS, which is widely known as endoplasmic reticulum and mitochondria visualization agent only.

Compared to most commercially available dyes, ABDS is cheaper, so it will be useful for biological laboratories that often face the need for a simple and inexpensive method to visualize living cells [26], as well as DIY biological devices such as smartphone-based microscopes [27]. For example, the typical price of the DiBAC dye is $70, while cost of red permanent marker is $1.5–$3. According to our calculations (see Sec. 3.7), one permanent marker lasts for approximately 100,000 cell staining cycles, which is sufficient to meet the basic needs of small laboratory groups. In addition, purchasing a red marker eliminates delivery issues and other bureaucratic red tape. We believe that ABDS will optimize the research activities of most biological laboratories, which will stimulate progress in biology, cytology, bioelectronics, and medicine.

## 2. Materials and Methods

### 2.1. ABDS preparation

To prepare ABDS, the 96% ethanol, a glass slide, phosphate buffered saline (PBS), and a permanent red marker (like red Edding E-140S) are required. The detailed preparation protocol is as follows (Figure 1):

1. In order to unify the amount of ink used for the dye, draw an open square of size 4 × 4 mm^2^ on a glass slide [Figure 1(a-d)] or any other pattern. The length of the line should be between 20 and 50 mm. We recommend using a special 3D-printed stencil, stl-file of which can be obtained by request or download from link Ref. [28].
2. Next, dissolve the depicted pattern with 50 *μ*l of 96% ethanol [Figure 1(e-f)].
3. After this, it is necessary to dilute 5 *μ*l of the resulting ethanol solution in 1 ml of PBS [Figure 1(g-h)]. To accurately control the ABDS concentration, an absorbance value of 0.05 at 527 nm for a 10 mm path can be used. For absorbance measurement, we have used polystyrol cuvettes (Sarstedt, Germany).

**Figure 1:**
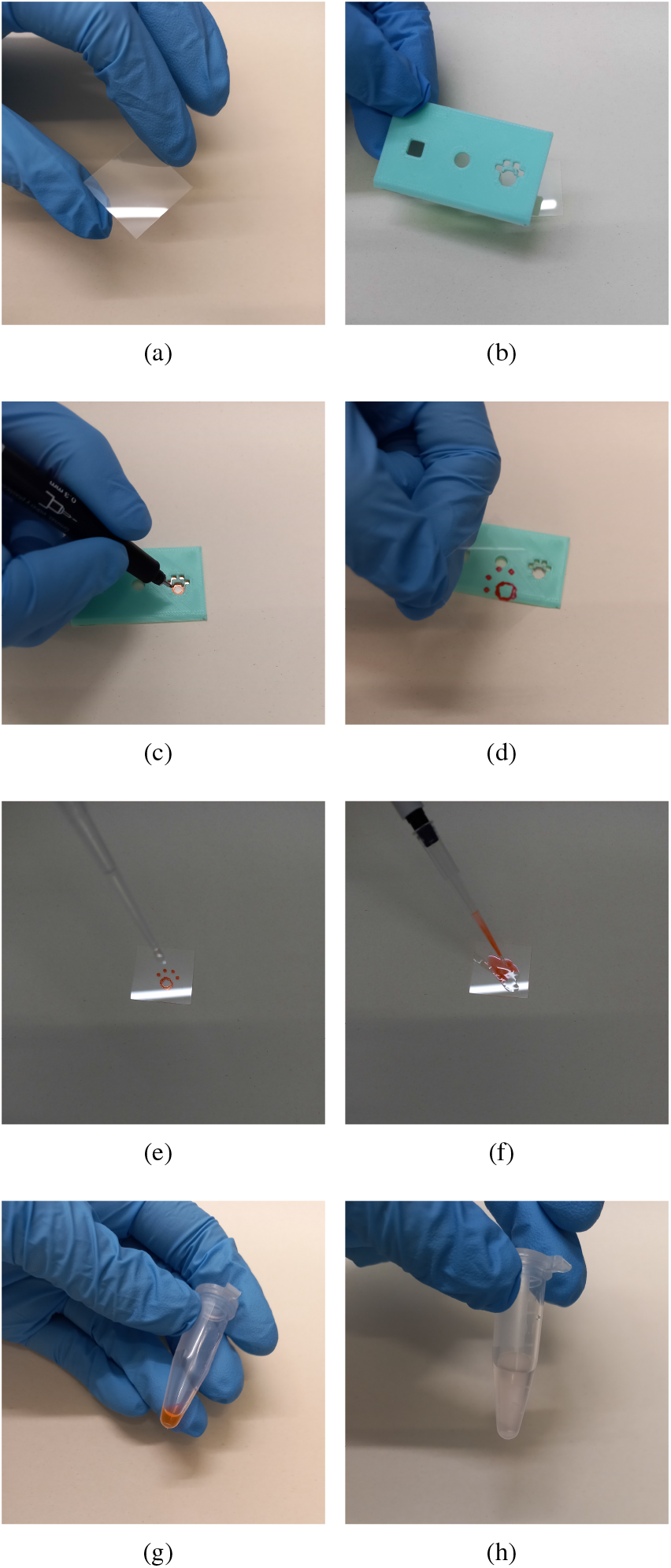
ABDS preparation protocol: (a) – glass slide, (b) – 3D printable spencil, (c) and (d) – drawing picture with red permanent marker, (e) and (f) – dissolving the picture with 50 *µ*l of 96% ethanol, (g) and (h) – dilution 5 *µ*l of the resulting ethanol solution in 1 ml of PBS.

Preliminary we have tested different types of markers (Figure 2) including red premanent markers Edding E140S (Edding International GmbH, Japan), Brauberg Classic (Brauberg, Russia), Crown Multi F-800W (Crown, USA) and Centropen 2836 (Centropen, Czech Republic), black, green, and blue permanent Edding E140S, red white-board marker Crown Multi WB-505 (Crown, Korea), and yellow and pink highlighters Edding 345 (Edding International GmbH, Japan). We found that bright fluorescence of cells was observed only when using red permanent markers from various manufacturers and the results were reproducible with all of them, suggesting that the red permanent marker ink uses the same active ingredient. So in our work we did not face the problem of markers fluorescence depending on the manufacturer mentioned in Ref. [19]. The green and blue markers from Edding do not provide a staining effect on cells. The ABDS-like dyes, prepared from pink and yellow highlighter pens, demonstrate fluorescence effect significantly lower than red permanent markers. The whiteboard markers and black marker from Edding do not demonstrate the fluorescence at all.

**Figure 2:**
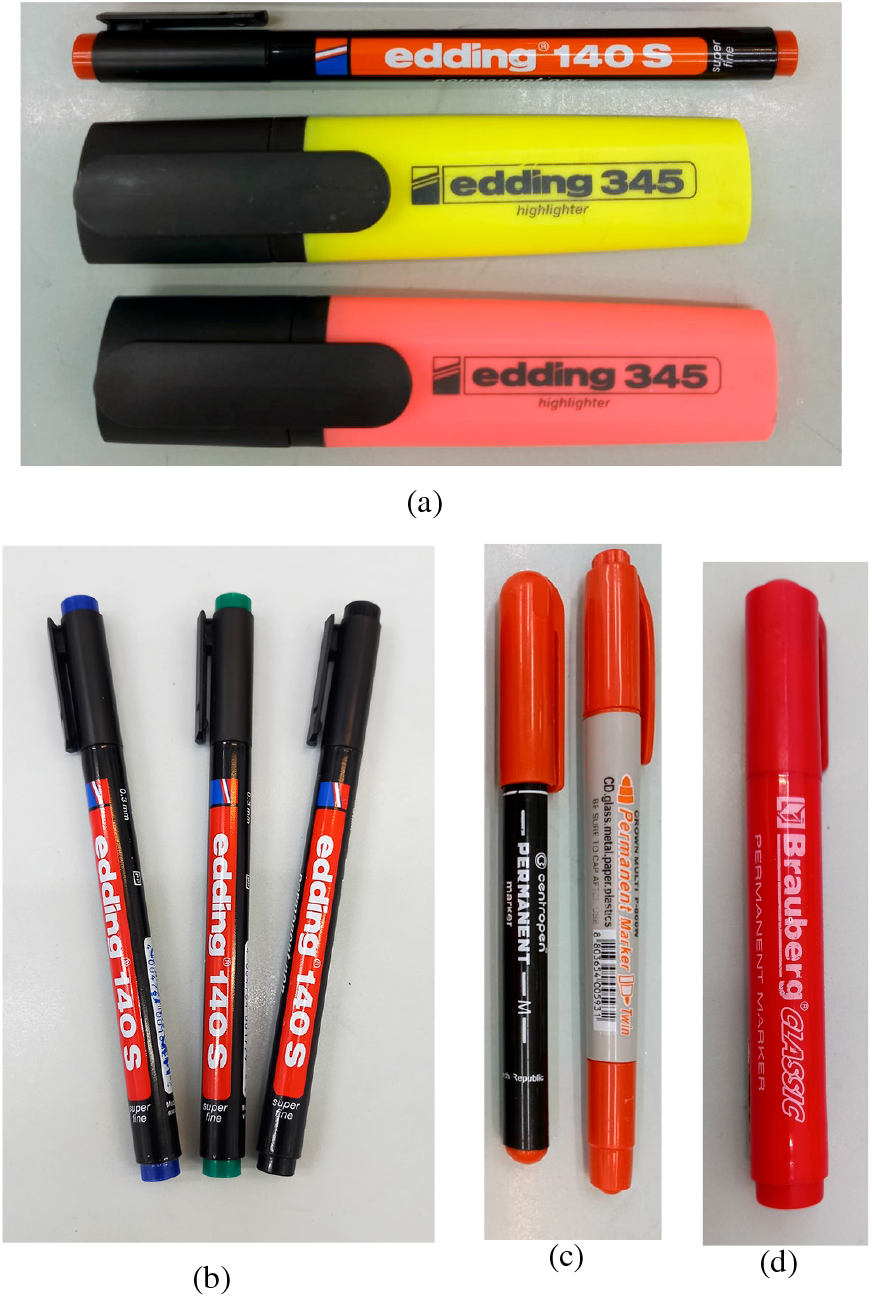
Markers types that have tested in this study.

Therefore, for ABDS preparation it is very important to use red and permanent markers, because inks with other colors and inks for whiteboard markers do not provide a fluorescence staining effect. This can be the result of the absence of the Rhodamine 6G component in them, which we assume provides a staining feature in the cells (see Sec. 3.2). In this study, we have mostly used red Edding E-140S marker with linewidth of 0.3 mm.

The ABDS dye is resistant to aggregation. However, for ABDS storage we recommend using glass tubes or using plastic tubes, but for no longer than 1 hour, since the dye tends to be absorbed into plastic.

### 2.2. 3D printable stencil

Since the initial concentration of the marker ink is typically unknown, we had to create a special stencil to control the concentration of ABDS, which has a square hole 5×5 mm^2^, a circle hole with 5 mm diameter and a cat paw hole, which we used for ABDS preparing (stl-file can be downloaded from link [28]). Also, the stencil has a rule cut for making an arbitrary length market ink line. The stencil is used for creating ABDS with reproducible concentration, *i*.*e*. it allows us to control marker ink amount by marker line length (for square hole the line will be have approximately 20 mm length, for circle hole it will be 5*π* mm, arbitrary length can labeled on the rule cut). The stencil was developed in Blender 4.3 software and printed from PLA plastic (Guangzhou Zhiwen Technology, China) on the Flying Bear S1 3D printer (Zhejiang FlyingBear 3D Technology Co., Ltd, China).

### 2.3. Optical spectroscopy study

In order to estimate the ABDS composition, we have measured its absorption, emission, and Raman spectra. The absorption was measured on the Evolution 300 UV-VIS spectrophotometer (ThermoFisher Scientific, USA) in the 400-700 nm range. The emission spectra were collected using a Chirascan spectrophotometer (Applied Photophysics, UK) using an excitation wavelengths of 527 nm (ABDS peak absorption), 532 nm, 488 nm, 365nm, and 639 nm (typical wavelengths used in fluorescence microscopy). The Raman spectra were obtained with the LabRam HR800 setup (Horiba, France) using IR laser 784.19 nm.

### 2.4. Cell cultures

In this study, we have used HeLa cells, obtained from the Bank of cell cultures of the Institute of Cytology of the Russian Academy of Sciences. For cell cultivation, we used the DMEM medium (Biolot, Russia). A 10% fetal bovine serum (HyClone, USA) and antibiotic gentamicin (Biolot, Russia) were added to cells medium before incubation, which was performed at 37°C and 5% CO_2_.

### 2.5. Intravital staining of the cells

Intravital cells staining with ABDS was carried out as follows. First, it is necessary to wash the cells from the culture medium with PBS at least twice. Then a PBS-solution of ABDS needs to be added to the cells (the preparation protocol is presented in Sec. 2.1), after which cells should be incubated with the ABDS-dye solution for 10 minutes at 37°C. Finally, the cells should be washed again with PBS twice. For immediate cells optical research, we recommend using optically transparent PBS as cells medium. Also, it is possible to incubate stained cells in culturing medium if it is planned to study them over several days. In this study, we have used analogical protocols for cells R6G, DiBAC and Deep Red staining.

### 2.6. Microscopy and third-party cell dyes

For ABDS staining examination we have used the Leica DM 4000 B fluorescence microscope equipped with EL6000 lamp (Leica, Germany, A, I3, and N2.1 cubes [29]) and Zeiss Observer.Z1 confocal microscope (Zeiss, Germany) equipped with fluorescence lamp HXP 120 (LEj JENA GmbH, Germany) and pumping lasers source with wavelengths 639 nm, 532/561 nm, and 488 nm (Zeiss Laser-modul TIRF, Germany). The Zeiss photographs were processed using AxioVision (SE64 Rel. 4.9), since the data obtained from the Zeiss microscope contains a larger than 256 number of tones (4096), which does not allow them to be processed using Adobe Photoshop (version 26.2.0), like photographs from a Leica microscope. The images were collected as 2D photographs and as 3D z-stack tomography. For high-resolution confocal pictures, immersion oil was used. As a sample for comparison, we have used the low-cost commercial dye DiBAC and moderate cost dye Deep Red Cell Mask (ThermoFisher Scientific, USA). The photographs are presented in monochrome or pseudo-color modes. Since ABDS composition demonstrates presence of Rhodamine 6G fluorophore (Sec. 3.2), we have utilized Rhodamine 6G from Sigma Aldrich (USA) as reference dye.

In our experiments with Zeiss lasers, we used 50% power for 532 nm and 75% power for 488 nm (standard fluorescence mode settings for the laser module used). Due to the difficulty of directly measuring laser power, we measured the photocurrent generated in a BPW20RF photodiode (Vishay Semiconductors, USA) when it was irradiated through an 100× objective lens by pump lasers. The resulting current was approximately 50 *μ*A. In this study, for cells R6G, DiBAC and Deep Red staining we have used protocol similar to presented one in Sec. 2.5.

### 2.7. Viability test

The viability test of the ABDS was performed using the cell BD FACSCanto Flow Cytometer (BD Bioscience, USA) and DAPI dye (dead cells nuclei visualization, ThermoFisher Scientific, USA) for non-alive cell detection. To monitor cells viability during the microscope study, we have also used the DAPI dye. All microscope photographs were made in the optically transparent non-fluorescence phosphate buffered saline (PBS, RosMedBio, Russia). For the MTS assay, which reflects the viability and proliferative activity of HeLa cells, the MTS homogenous reagent (Himedia, India) and Multiskan 60 Microplate Spectrophotometer (Thermo fisher scientific, USA) were used.

### 2.8. Phototoxicity test

To conduct the cytotoxicity test, we used the following experimental design. Cells incubated on thin-bottomed Petri dishes were stained with ABDS for one sample type and DiBAC for the other. Control cells were not stained. DAPI was added to the medium for each sample as an indicator of cell death. Each sample was then continuously irradiated with a 532 nm or 488 nm laser through 10x or 100x objectives on a Zeiss Observer.Z1 microscope. During irradiation, the cell was photographed once per minute in the brightfield, laser fluorescence, and ultraviolet (UV) fluorescence channels. To prevent cells damage due to UV radiation, the time for obtaining photographs in UV mode was limited to 5 s.

## 3. Results and discussion

### 3.1. ABDS fluorescence study using microscopes

To study ABDS cells staining feature, we have provided a wide range of experiments with fluorescence microscopy. At first, we have examined an existence of the ABDS fluorescence in HeLa cells with a Leica DM 4000 B fluorescence microscope (Leica, Germany). The results obtained for cubes I3, N2.1 and A, as well as a reference bright-field image of the cells are shown in Figure 3(a), which clearly demonstrates that ABDS successfully stains the cells and allows to obtain their high-quality images. Photographs of cells in Figure 3(a) were processed with the highest image clarity using the “curves” tool in Adobe Photoshop [shape of the curves for each of the photographs is shown in bottom panels in Figure 3(a)]. It can be seen that the ABDS photograph obtained with cube N2.1 (green light pumping, red light emission) has the best image quality, the photograph obtained with cube I3 (blue light pumping, green light emission) has less contrast, and cube A with UV pumping demonstrates very weak fluorescence signal. These results show that the green light range is preferable for ABDS pumping, which is confirmed by fluorescence spectra analysis [Figure 4(a)]. It also follows from Figure 3(a) and Figure 4(a) that ABDS is compatible with UV-pumped blue dyes such as DAPI since ABDS fluorescence is very weak when 365 nm excitation is used. Moreover, as can be seen from the optical spectra in Figure 4(a) and the live-cell photographs in Figure 3, ABDS does not require specific 488 nm-only or 532 nm-only pump sources, making its use more accessible to most researchers. On the other hand, the problem of ABDS fluorescence overlap with other fluorophores that are excited only at 488 nm or 532 nm can be easily resolved by numerical subtraction of images, since we observed that the ratio between ABDS fluorescence excited at 532 nm and 488 nm is constant in time and space when other excitation parameters are fixed. We exploited this effect in Sec. 3.3 to separate the fluorescence signals of DiBAC and ABDS.

**Figure 3:**
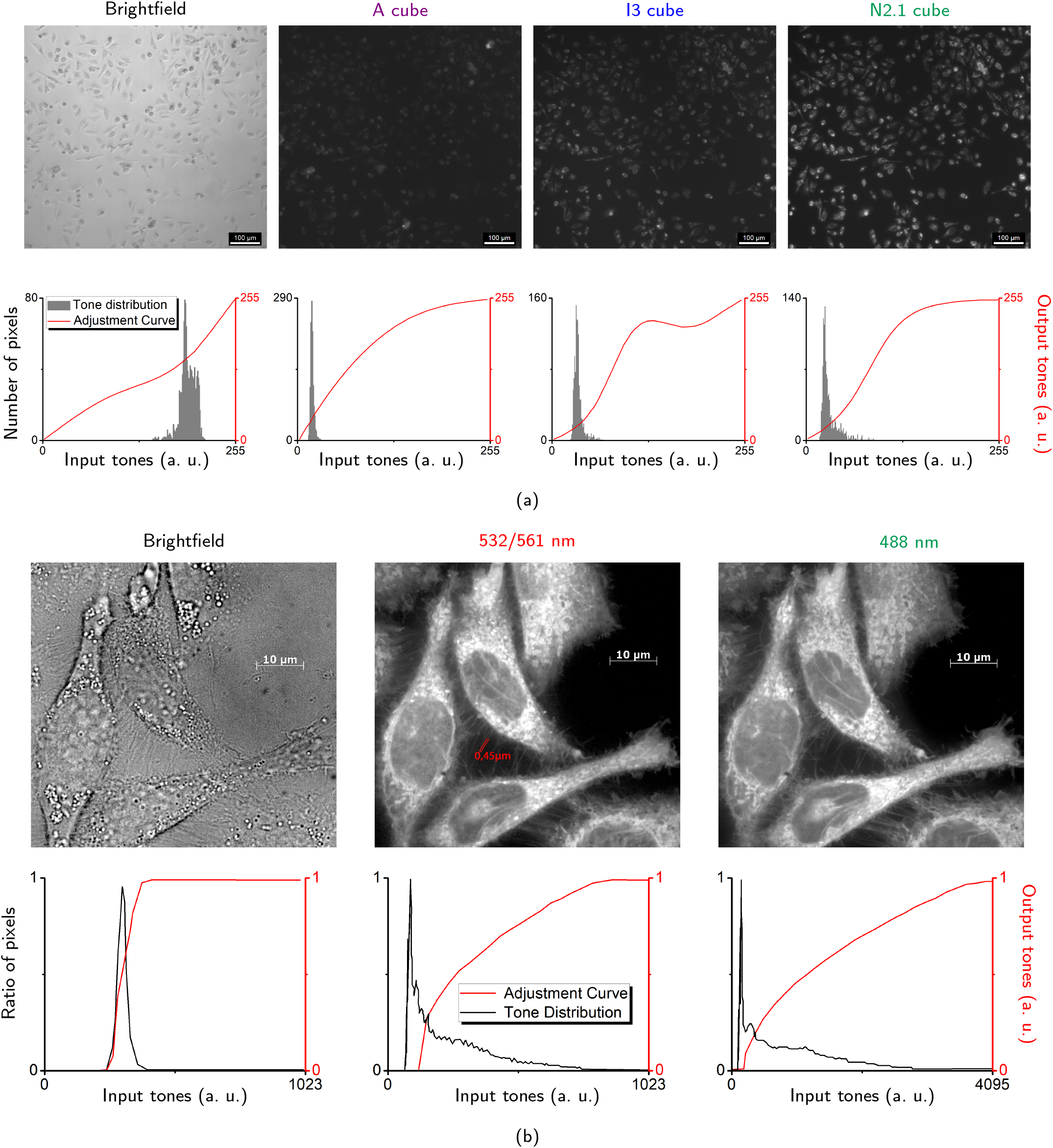
ABDS fluorescence in HeLa cells. (a) Photographs obtained with Leica DM 4000 B fluorescence microscope. With this microscope cells were studied using brightfield mode, and fluorescence mode with I3, N2.1, and A cubes. The photographs’ tones were processed by Adobe Photoshop. (b) Photographs obtained with Zeiss Observer.Z1 confocal microscope. With this microscope cells were studied using brightfield, and in fluorescence mode with lasers 532/561 nm and 488 nm. The photographs were processed by AxioVision (tuning curves are presented at bottom panels).] The obtained data indicates that it is possible to study cells’ membrane structure with submicron resolution using ABDS (cmp. with Figures 5 and 6). The difference in the tone range in histograms for lasers can be due to different lasing light intensity, which is typically unknown and very difficult to measure.

**Figure 4:**
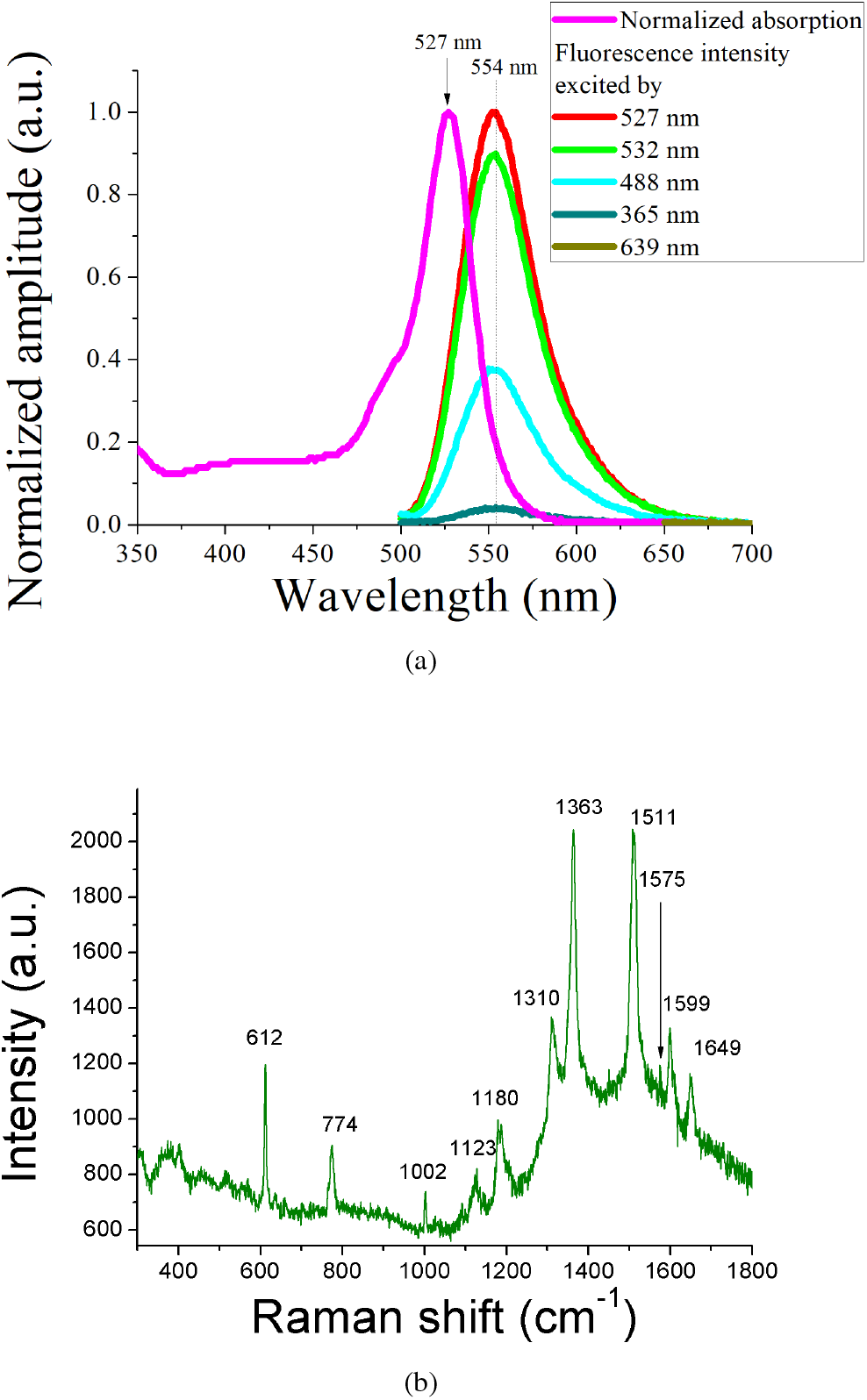
Optical spectroscopy study of ABDS: (a) – absorption and emission ABDS spectra, (b) – Raman spectrum of ABDS. The peaks signature is close to Rhodamine 6G [34]. Emission spectra were normalized on fluorescence intensity maximum for 527 nm excitation. The short wavelength vertical offset in ABDS absorption spectrum is not a baseline artifact, since its existence allows for excitation of ABDS by a high-energy photons, such as 488 nm. The same offset is also observed in the R6G spectrum Figure 14(a).

The more striking result of the ABDS HeLa staining isshown in Figure 3(b), where obtained with Zeiss microscope confocal images of the HeLa cells treated with ABDS are presented. These photographs demonstrate the submicron resolution of the ABDS dye, which allows visualization of resolution of the ABDS dye, which allows visualization of not only membrane topology but also cytoskeletal structures like filopodia (see Sec. 3.3). Similar to the Leica data, the Zeiss raw images were adjusted with AxioVision software for better clarity. From the adjusted images in Figure 3(b) it can be concluded that pumping in the green and blue light lasers are suitable for ABDS excitation. In addition, we also observed no ABDS signal with usage of red laser pumping 639 nm, which makes ABDS compatible with red fluorescence dyes like Deep Red Cell Mask. This statement is also supported by Figure 4(a), where no fluorescence was detected when ABDS was pumped by a 639 nm laser, since ABDS does not absorb light at this wavelength.

### 3.2. ABDS absorption, emission, and Raman spectra

Since the exact composition of the Edding 140-S permanent red marker ink is unknown and most likely is a trade secret, we attempted to reverse engineer the fluorescent substance in the ink. In order to determine which substance fluoresces in the composition of ABDS, we examined it using absorption, emission, and Raman spectroscopy. The results are shown in Figure 4. Since fluorescent staining could be carried out not only with the help of the marker Edding 140-S, but also with red fluorescent markers from other companies, we concluded that the substance used as a dye is widely used. Search in various literary sources led us to the assumption that the coloring pigment of red alcohol ink or one of its components is the substance Rhodamine 6G [30, 31]. At the same time, the absorption [32], emission [33], and Raman spectra are known for Rhodamine 6G [34], and coincide with the ABDS spectra [compare Figures 4(a) and 14(a)]. Thus, we can assume that Rhodamine 6G is highly probable present in the composition of the ABDS dye. Possible additional peaks in the Raman spectra of the ABDS may mean that the ink of the permanent red marker also contains other substances, which we, however, had not identified yet.

We have measured the emission spectra pumping ABDS at the its absorption maximum 527 nm, and at levels 365 nm, 488 nm, 532 nm, and 639 nm, which are used in the fluorescence microscopy. As a result, we observed no fluorescence while 639-nm pumping is used, and the 50-nm width emission peaks at 554 nm, when other pumping levels are used. It should be noted that these peaks of ABDS fluorescence are close to to the maximum sensitivity of the human eye [35] and are therefore very clearly visible under a microscope (Figure 3). Surprisingly, no other peaks were detected in the fluorescence spectrum, that means that ABDS does not change color with respect to excitation wavelength change and ABDS color images obtained by fluorescence cubes just cut off fluorescence signal associated with 554 nm peaks. The absence of the other peaks can be explained by the energy diagram proposed in the work by [36]. Indeed, let assume that Rhodamine 6G in ABDS exhibits a radiative transition between the *S*_1_ → *S*_0_ states, while the others are nonradiative. As can be seen from the absorption spectra Figure 4(a) and 14(a), ABDS and Rhodamine 6G can slightly absorb high-energy photons, which correspond to higher energy states. However, since they are nonradiative states, the system relaxes nonradiatively to the *S*_1_ state and then emits a photon by relaxation to the *S*_0_ state. Therefore, the peaks position and their widths are independent of the pump wavelength if it is lower than 527 nm, while the peak intensity decreases.

### 3.3. ABDS labeling regions. Proving of the membrane staining property

As shown in section 3.2, ABDS is highly likely to contains Rhodamine 6G as a component, so it should stain mitochondria and the endoplasmic reticulum [37, 38, 39]. However, as shown in Figure 3(b), ABDS allows visualization of filopodia-like 450-nm structures that are typically associated with the cell membrane. This observation, along with previously published studies measuring the integral membrane potential of the cell using Rhodamine 6G [40], leads us to hypothesize that ABDS/R6G also stains the cell membrane. To prove this, we performed series of experiments in which HeLa cells were stained with ABDS and Rhodamine 6G and with commercial dyes classified as membrane stainers, namely Deep Red Cell Mask and DiBAC (ThermoFisher, USA). The obtained results are presented in Figures 5, 6, 7, and 8.

**Figure 5:**
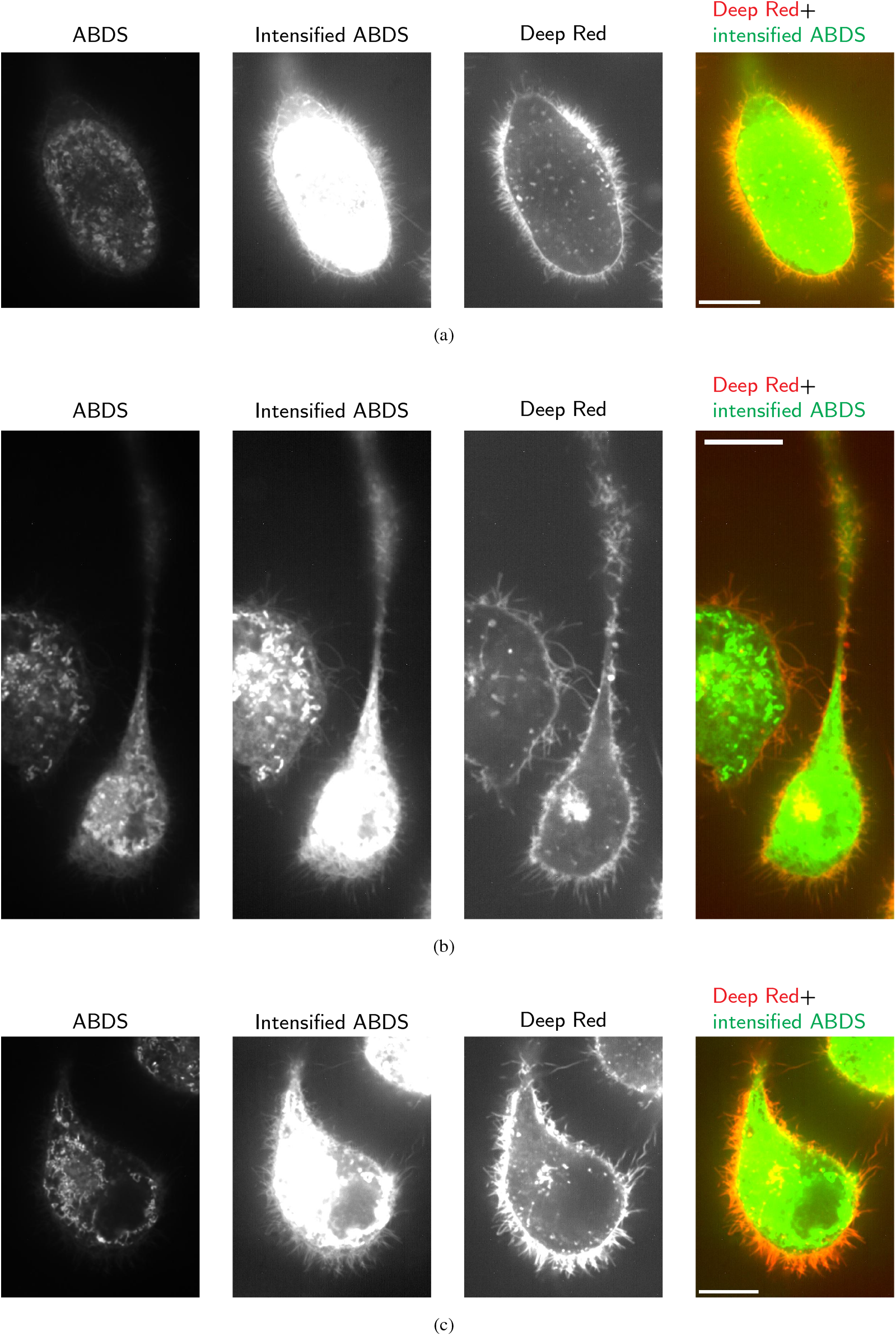
Proof of the cytoplasmic membrane staining property of the ABDS. One can see that intensified ABDS signal contains the cytoplasmic membrane images. Intensification was provided by gamma-correction and scaling. Scale bar corresponds to 10 *µ*m. The video compilation is available at https://www.youtube.com/shorts/NGaPd9hzy4k

**Figure 6:**
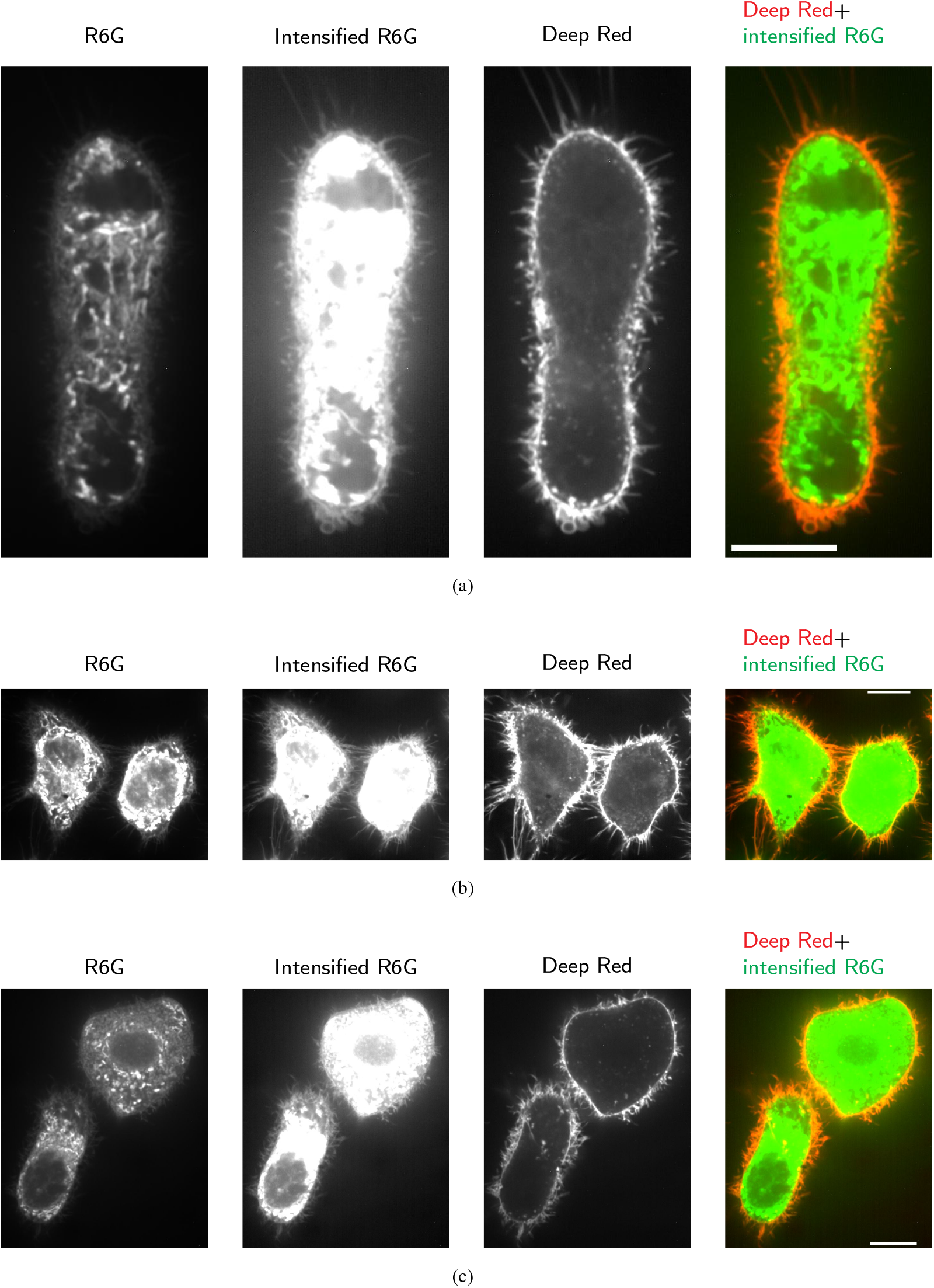
Proof of the cytoplasmic membrane staining property of the Rhodamine 6G. Similar to Figure 5, we can see that intensified Rhodamine 6G image contains the signal from the membrane. Intensification is produced similar to the Figure 5 data. Scale bar corresponds to 10 *µ*m. The video compilation is available at https://www.youtube.com/shorts/NGaPd9hzy4k

**Figure 7:**
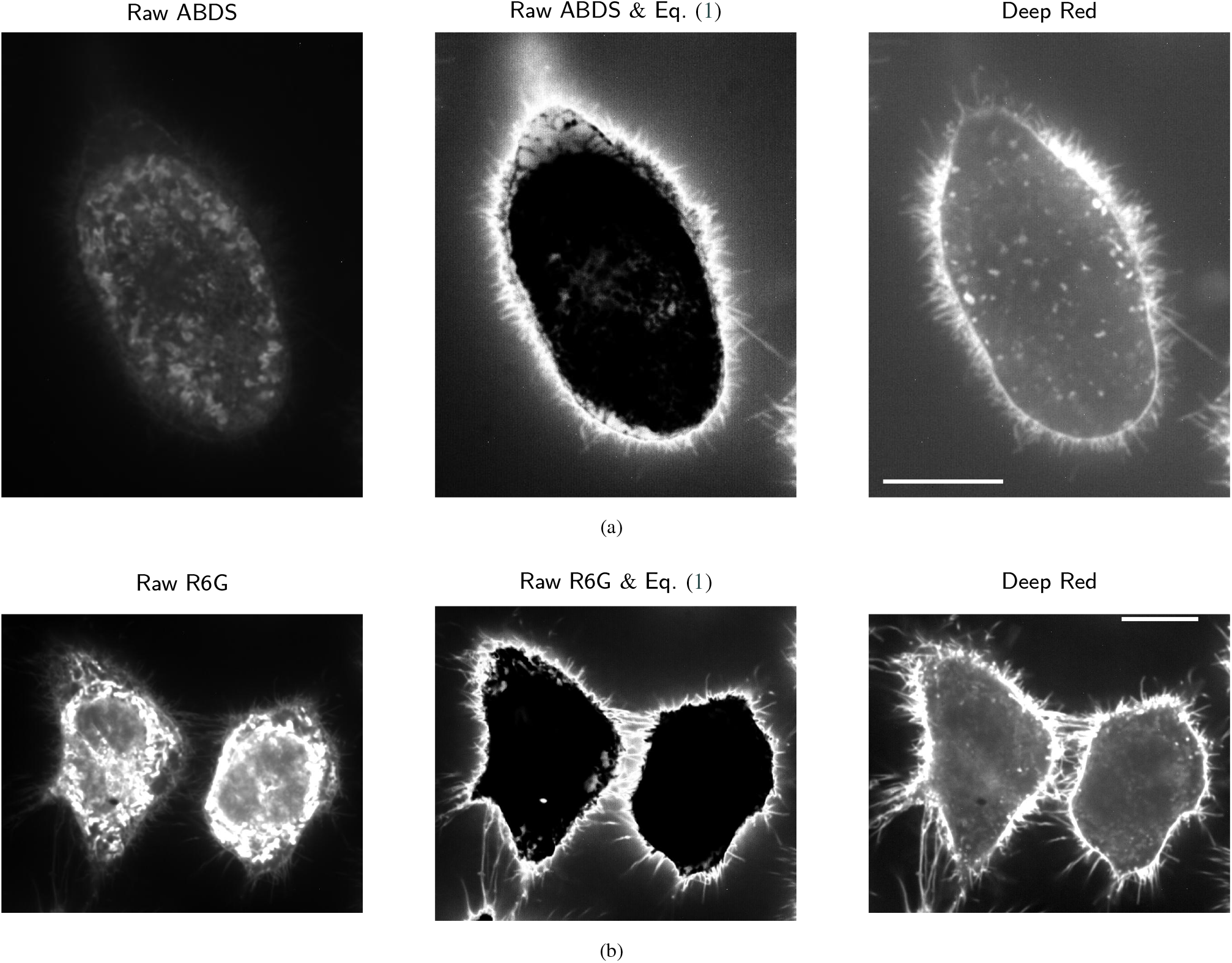
Extraction of cell membrane signal from ABDS (a) and R6G (b) images using a simple thresholding technique based on applying a Gaussian bell-shaped function Eq. (1) to the raw images. This procedure clearly suppresses the endoplasmic reticulum signal and enhances the membrane signal intensity, further confirming the ability of ABDS/R6G to stain the cell membrane. Scale bars corresponds to 10 *µ*m.

**Figure 8:**
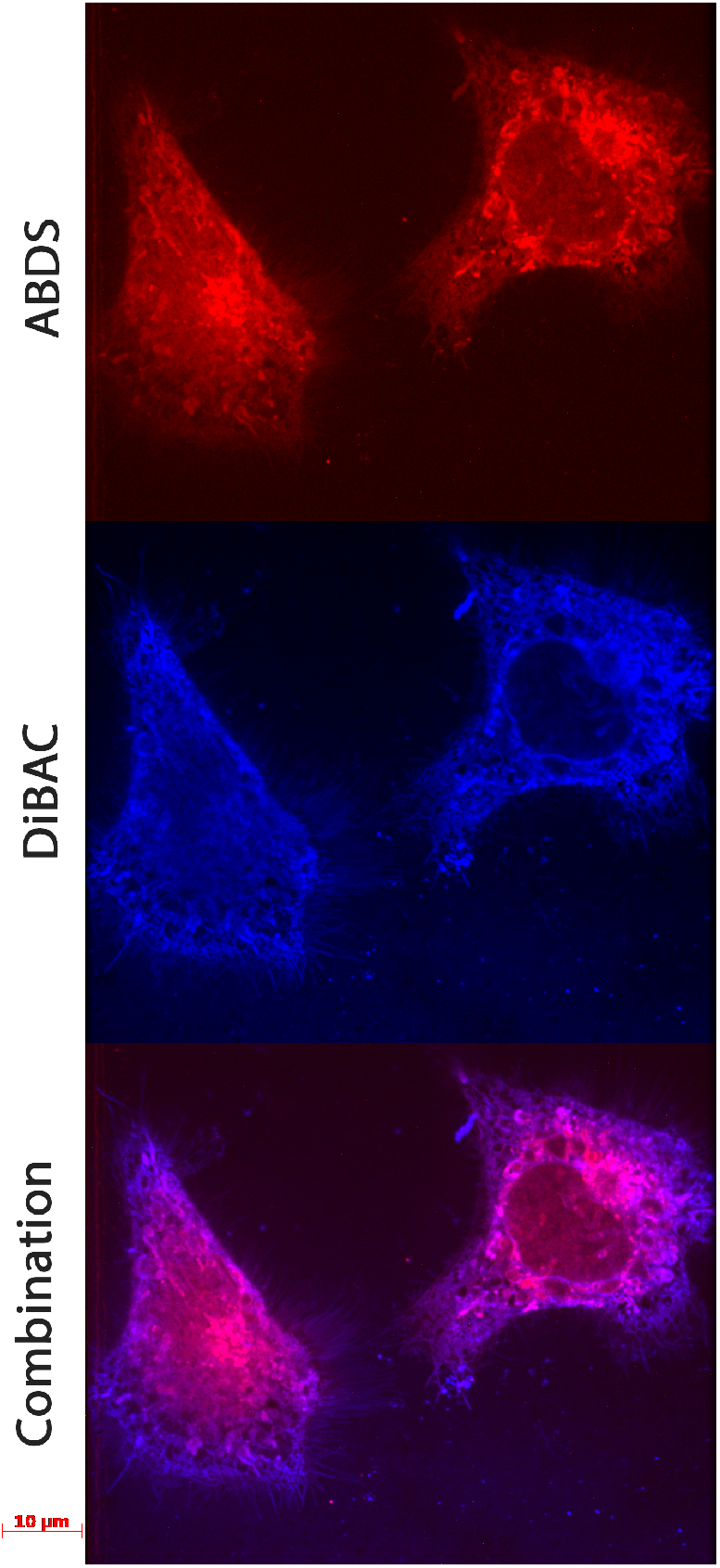
Comparison of the ABDS image with the DiBAC signal. To suppress the ABDS signal in the DiBAC image, we detected cells in the probe for which the DiBAC intensity is higher than the ABDS intensity, the amplitude of which we monitored using the 532 nm pump channel, where DiBAC fluorescence was absent (the ABDS fluorescence intensities in the 532 nm and 488 nm channels are related as 1:2 for the pumping settings we used). The resulting image clearly demonstrates that the ABDS fluorophore is localized predominantly in the same location as DiBAC, which allows us to conclude that these dyes stain the same areas of the cell.

The most groundbreaking outcome of such head-to-head experiments can be done by comparing ABDS/R6G and the pure membrane dye Deep Red Cell Mask, as shown in Figures 5 and 6. From these photographs it is evident that the intensified ABDS/R6G image also includes the cell membrane signal. The membrane signal intensity of ABDS/R6G is lower than the intracellular fluorescence intensity, which we attribute to the endoplasmic reticulum, and this is perhaps why the effect of membrane staining with Rhoadmin 6G has not yet been detected. Surprisingly, in some cases, this low membrane signal amplitude allows us to isolate its image from ABDS/R6G images using simple double “soft” thresholding. For example, by processing the ABDS/R6G signal intensity *I* with a bell-shaped Gaussian function

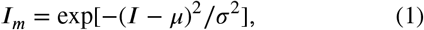

where *μ* corresponds to the average membrane signal intensity and *σ* is the range of membrane signal intensities, we can enhance the low-intensity membrane signal *I*_*m*_ and suppress the background signal and the high-intensity endoplasmic reticulum signal, as shown in Figure 7. Thus, the presented data proved the ability of ABDS/R6G to stain the cytoplasmic membrane. More accurate and versatile membrane signal extraction from ABDS/R6G data can be achieved using AI-based image processing, which will be discussed in our future articles. The alpha-versions of the neuron nets for membrane extraction can be found at our Github repository [41]. To our knowledge, only one recent study [42] claims that R6G stains cell membranes in yeast. However, the authors of this paper did not prove this claim or examine the quality of R6G membrane staining or its application in eukaryotic cell research, which is what we did in the present study.

To experimentally reference ABDS labeling regions, we also compared ABDS with DiBAC, whose fluorescence images appear similar to ABDS. However, these experiments required some additional image processing tricks, since the fluorescence signals of DiBAC and ABDS overlap when using 488 nm excitation [Figure 4(a) and Figure 14(b)]. To address this issue, we found that the ratio between the fluorescence intensities of ABDS excited at 488 nm (*I*_488_) and 532 nm (*I*_532_) is constant in space and time for fixed excitation and fluorescence acquisition settings. This means that ABDS images acquired at 488 nm and 532 nm can be reconstructed from each other. For the data acquisition settings that we used (50% of the 532 laser power and 75% of the 488 laser power) this ratio *I*_488_/*I*_532_ ≃ 2. Using this feature of ABDS, we prepared experimental conditions in which the DiBAC fluorescence intensity in the 488 nm excitation channel became approximately 30 times higher than the ABDS intensity in this channel, which is monitored by the ABDS intensity in the 532 nm excitation channel, where no DiBAC fluorescence exists, since DiBAC does not adsorb light at this wavelength [Figure 14(b)]. This allows us to neglect the ABDS fluorescence signal in the 488 nm excitation channel and interpret it only as a DiBAC signal, while ABDS localization can be achieved by the 532 nm excitation channel. The results are shown in Figure 8. It is evident that ABDS predominantly stains the same area as DiBAC, from which we can conclude that these dyes are specific for the same cellular regions.

### 3.4. Cytotoxicity and phototoxicity tests

In order to determine the cytotoxicity of ABDS, a flow cytometric study and MTS test were conducted. The dye-containing samples were HeLa cells stained with ABDS according to the protocol given in section 2.5. HeLa cells unstained by ABDS were used as a control. For MTS assay cells were seeded in a 96-well plate and stained with ABDS at concentrations of 0.1×, 1×, 10×, 50×, and 100× from the standard ABDS concentration (see Sec. 2.1) for 24 hours in five wells for each concentration. To compare the cytotoxicity of ABDS 1× with commercial dyes, cells treated with DiBAC were used. We used 1 *μ*l of DiBAC per 1 ml of PBS solution (absorbance at 488 nm for 10 mm path was 0.25). The results of the study are shown in Figure 9. As can be seen in the figure, ABDS is not cytotoxic in 0.1× and 1× concentration, since the number of dead cells in stained samples is statistically indistinguishable from the number of dead cells in control samples. Moreover, even for 10× concentration about 80% of the cells remain alive with respect to control. Only in concentration 100× the cells viability became very low, which can be connected with ethanol presence in the ABDS staining solution. Finally, no significant difference between viability of the cells treated with 1× ABDS and DiBAC was observed, which means that ABDS cytotoxity is similar to commercial dyes.

**Figure 9:**
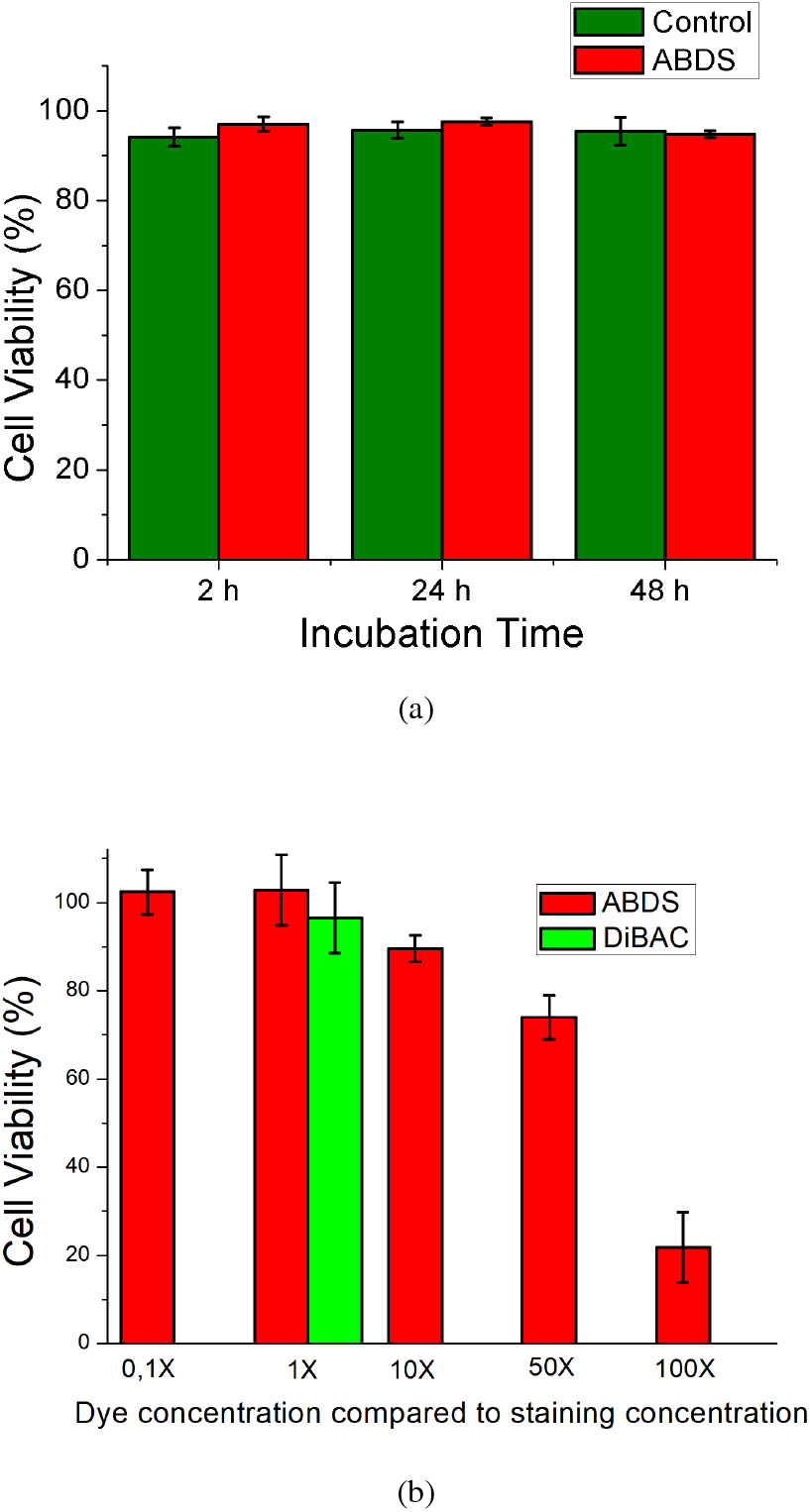
Cytotoxicity test. (a) Using cell flow cytometry. When processing the flow cytometry results, the number of surviving cells was compared with the initial number of living cells in the samples. The number of surviving cells in the control sample and stained by ABDS sample is statistically indistinguishable after 2, 24 and 48 hours, therefore, ABDS is not toxic to cells. Error bars are calculated as 95% confidence intervals. (b) Using MTS test. ABDS is not cytotoxic in 0.1×, 1× and 10× concentrations and it’s impact on cell viability for 1× concentration is close to impact of the DiBAC dye.

The qualitative phototoxicity test was provided using protocol from Sec. 2.8. As a result, we have found that neither ABDS, nor DiBAC do not have an affect on cells viability, which were irradiated by 532 nm and 488 nm lasers during 20 min using 10× objective. In contrast, for 100× objective we have observed that after 5-7 minutes irradiation cells stained by both dyes are dying, which is detected by DAPI fluorescence appearing, however control cells demonstrate such fluorescence effect only after irradiation during 12 min. Thus, we have determined that ABDS phototoxicity is the same as DiBACs.

### 3.5. Fluorescence photo-bleaching test

Various experiments with living cells, in particular in medicine and pharmacology areas, may require a long period of observation, including several days (a cell proliferation study [43], *in vitro* drug testing [44, 45], *etc*.), which requires the dye to remain stable throughout the observation period. In this context, one of the most interesting properties of ABDS is its ability to not fade for a long time and even increase fluoresce brightness for several minutes under constant laser pumping.

The problem of dyes’ fading is often a significant obstacle to recording experiments in time-lapse mode and z-stack scanning of cells. In both cases, the initial frames are clear and contrast, but over time the typical dyes begin to fade, and, if the study requires a large number of frames, near the end of the experiment, the photographs become more blurry and significantly less informative and representative. In contrast to typical dyes, ABDS is very stable for long pumping times, as shown by the results of the exposure stress test, which we provide for ABDS and the commercial intravital dye DiBAC. During this test, we compared the ability of ABDS and DiBAC to maintain fluorescence at an acceptable level over long-term experiments. The stress exposure test consisted of continuously irradiating cells stained by ABDS and cells stained by DiBAC with a Zeiss fluorescence microscopy pumping laser (see Sec. 2.6) with emission wavelength 488 nm. This wavelength excites the DiBAC dye, and we also used it to pump ABDS to ensure that the dyes were on equal terms. The cells were continuously illuminated by the laser emission for 20 min except time periods when photograph occurred. The photographs were taken with a 1-second exposure. The obtained results are presented in Figure 10. The photographs were processed using the settings “BestFit” of the program AxioVision. As can be seen from photographs sequences, over time, in about 10 minutes after exposure the fluorescence of DiBAC becomes less bright and, as a result, the pictures become less clear. This problem with fluorescence becomes especially significant when careful comparison of samples before and after the test treatment is necessary, since it is impossible to tell whether the changes are due to the experimental treatment or to dye burnout. In contrast, the dependence of the fluorescence of the ABDS on the exposure dose is not trivial. ABDS fluorescence initially increases for about 15 minutes (cmp. with Figure 11) and then the fluorescence becomes less bright. However, it should be noted that after 20 minutes of the experiment, the fluorescence did not drop below the initial level. This useful property of the ABDS dye can be utilized for many applications, for example to observe cell division, apoptosis process, de-adhesion study, *etc*.

**Figure 10:**
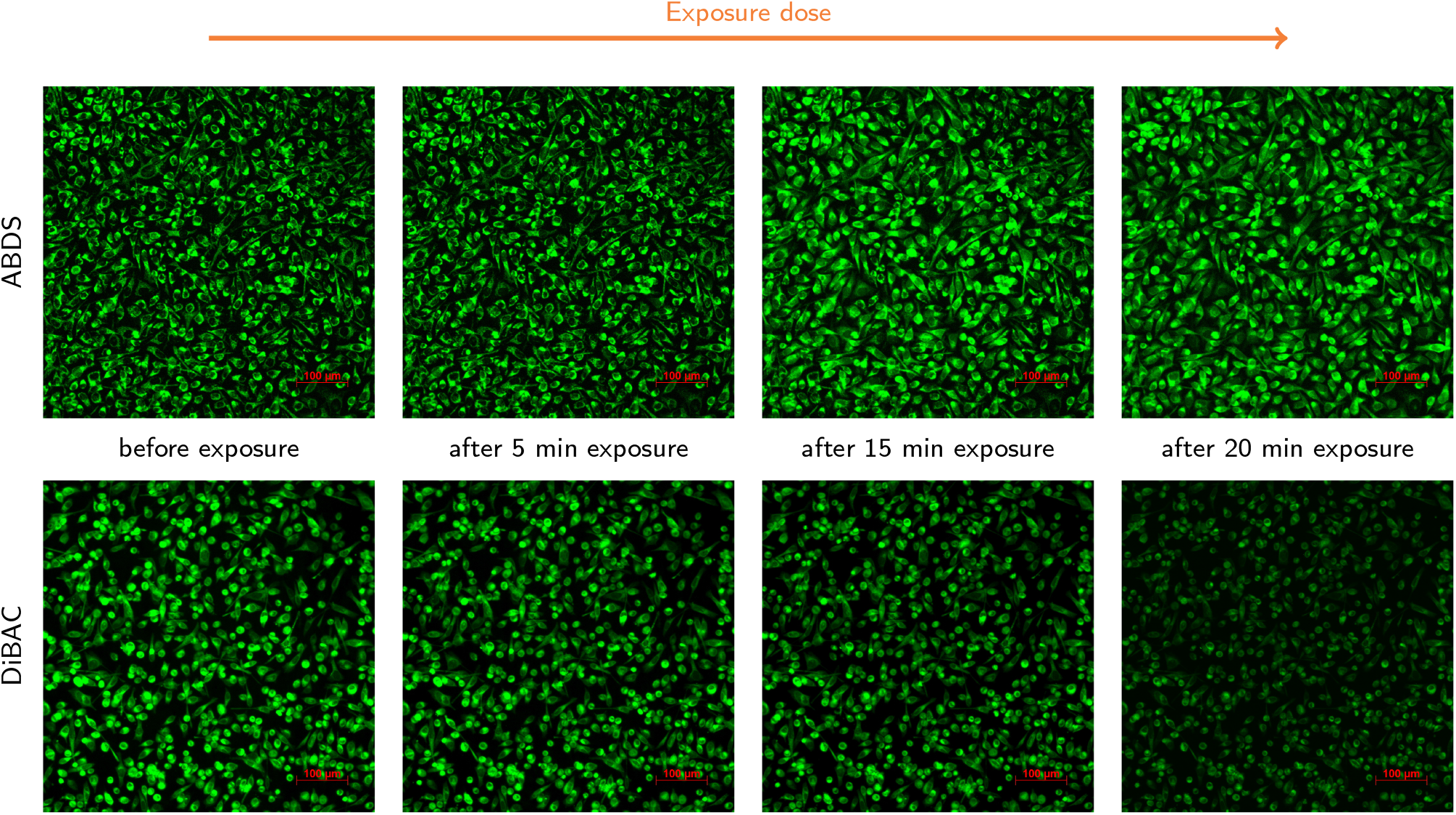
Exposure dose stress test for ABDS and DiBAC. It can be seen that after 15 minutes the fluorescence of the ABDS dye becomes brighter and then after 20 minutes it fades slightly. In contrast, fluorescence of the DiBAC dye just becomes less brighter with the time and, as a result, the photograph becomes less clear. The photographs were taken at 10X magnification with 488 nm laser on 75% of max power. Exposition for photographs was 1000 ms.

**Figure 11:**
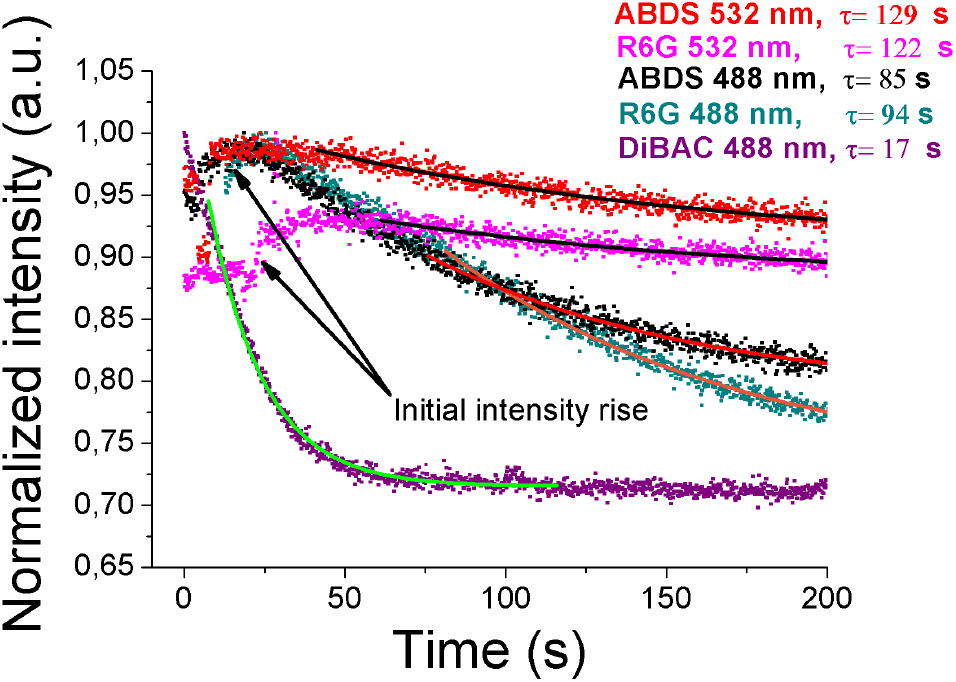
Photobleaching test for ABDS, Rhodamine 6G (R6G), and DiBAC. Fluorescence intensity dynamics obtained with continuous 532-nm and 488-nm laser pumping of the HeLa cells under 100x magnification using capturing signal from live camera “CSU-X1 Confocal Scanner Unit” (Yokogawa, Japan) by BandiCam software. Intensities are normalized by their maximum values. The scatter points correspond to experimental data, and solid lines corresponds to the intensity data I(t) data fitting to the empirical formula I(t)= *A* + *B* exp(−t/*τ*), where *τ* is the characteristic photo-bleaching time. The sample live capture of the ABDS and DiBAC photobleaching can be found at https://youtu.be/OPd_vfpFJ4k?si=-kGhi7guIkOrsFvL.

The similar dye burnout stress test for 100X magnification presented in Figure 11 demonstrates that DiBAC fading out within 20 s, which is insufficient even for aiming. In contrast, ABDS and Rhodamine 6G are stable for 1.5 minutes. The real-time comparison between ABDS and DiBAC photobleaching can be seen from video at link https://youtube/OPd_vfpFJ4k?si=-kGhi7guIkOrsFvL. Moreover, when 532-nm pumpimg is used, both ABDS and Rhodamine became stable for 2 minutes. Thus, ABDS can be used for capturing cells images in the cases when utilization of the convenient dyes is complicated.

The nature of the increasing ABDS intensity can be explained by quantum quenching phenomena [46]. Indeed, when the concentration of ABDS is high, then emitted photons from one fluorescence center have high probability to absorb in another, where it can relax in a nonradiative way. However, during the pumping process some fluorescence centers start to burn down, which decreases the overall dye concentration and thus decreases the probability of the nonradiative relaxation. The latter led to an increase of the total ABDS fluorescence intensity.

It should be noted that a qualitatively similar intensity increase effect was recently described in Ref. [47] for Rhodamine 6G, where the authors isolated a droplet of rhodamine using electrostatic levitation. In our study, we also isolated a small amount of ABDS/R6G, but for this purpose, we used simple biological cells. This observation further supports the idea that ABDS is based on rhodamine 6G. However, we also observe that the time evolution of ABDS fluorescence intensity, in contrast to the study in Ref. [47], has an asymmetric curve, and its decay can be successfully approximated by a single-exponential model, rather than the quadruple-exponential model proposed in Ref. [47] (see Figure 11).

### 3.6. High Detalization and z-stack tomography

Another advantage of ABDS is its ability to highlight focal contacts – the protein processes of adhesion cells, which they use to attach to the surface of the culture dish [48] [Figure 3(b), highlighted by 0.45 *μ*m marker]. Of the commercially available dyes, the Deep Red Cell Mask has the ability to provide a sufficiently detailed picture. A comparison of photographs obtained using a Zeiss Observer.Z1 confocal microscope for Deep Red Cell Mask and ABDS dyes is shown in Figure 5. As can be seen in these photographs, ABDS paints focal contacts as well as Deep Red Cell Mask, despite the price difference of more than a hundred times (see Sec. 3.7).

Moreover, the obvious application of the ABDS fluorescence stability and brightness feature (see Sec. 3.5) can be found in the living cells high-magnification confocal tomography. Figure 12 and Figure 13 show a comparison of z-stack images of the HeLa cells for ABDS and DiBAC. It can be seen that when staining cells with DiBAC, the dye burns out already when the first few slices are captured. This property of the dye to burn out is especially visible when displaying the results in the Cut-view mode, shown in Figure 13. Because of this DiBAC feature, 3D-tomography of cells with it at 100X magnification is ineffective. At the same time, ABDS does not burn out under the same shooting parameters, which in contrast to DiBAC-like dyes allows obtaining both clear Cut-view and detailed 3D tomography of cells.

**Figure 12:**
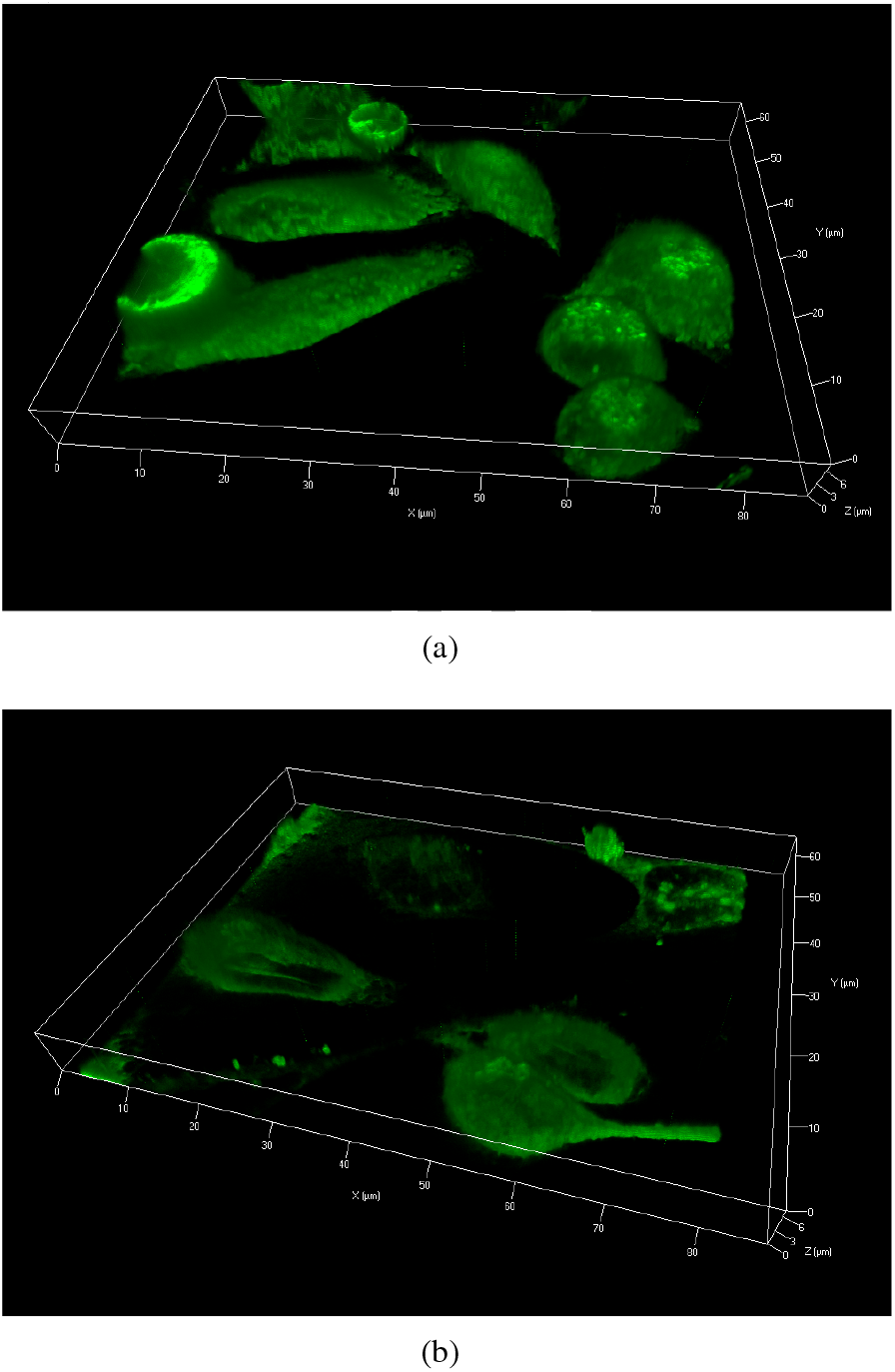
HeLa cells z-stack 3D tomography stained with ABDS (a) and DiBAC (b). It can be seen that DiBAC burns out during photographing the lower layers of the cell, while ABDS allows to construct a complete 3D-tomography of the cell. The exposure was 1000 ms per slice. Additional demos are available at https://www.youtube.com/watch?v=sPwg2KGtUz8 and https://www.youtube.com/watch?v=OPd_vfpFJ4k.

**Figure 13:**
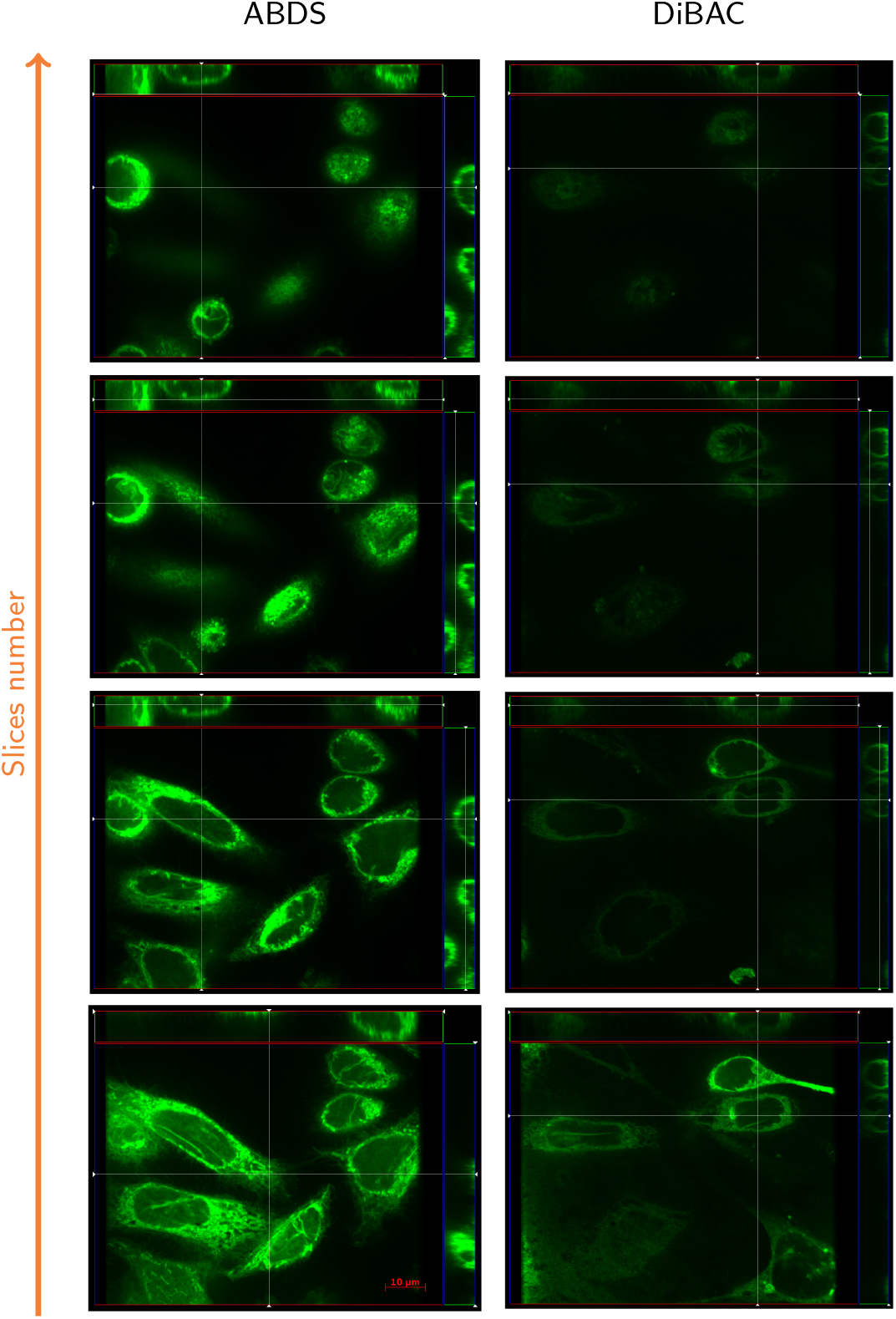
Cut-view of HeLa cells stained with ABDS and DiBAC respectively. It is evident that when moving from the lower slices to the upper ones, DiBAC burns out a lot, making a full z-stack of cells almost impossible [see Figure 12(b)]. At the same time, ABDS does not burn out in the same number of frames and allows for high-precision photographing of both the lower slices and the upper part of the cells [see Figure 12(a)]. This figure is consistent with the data presented in Figure 11. The signal-to-noise ratio (SNR) for ABDS for top and bottom slices can be estimated as 37 dB, while DiBAC SNR for bottom slice is 20 dB and 13 dB for top slice.

**Figure 14:**
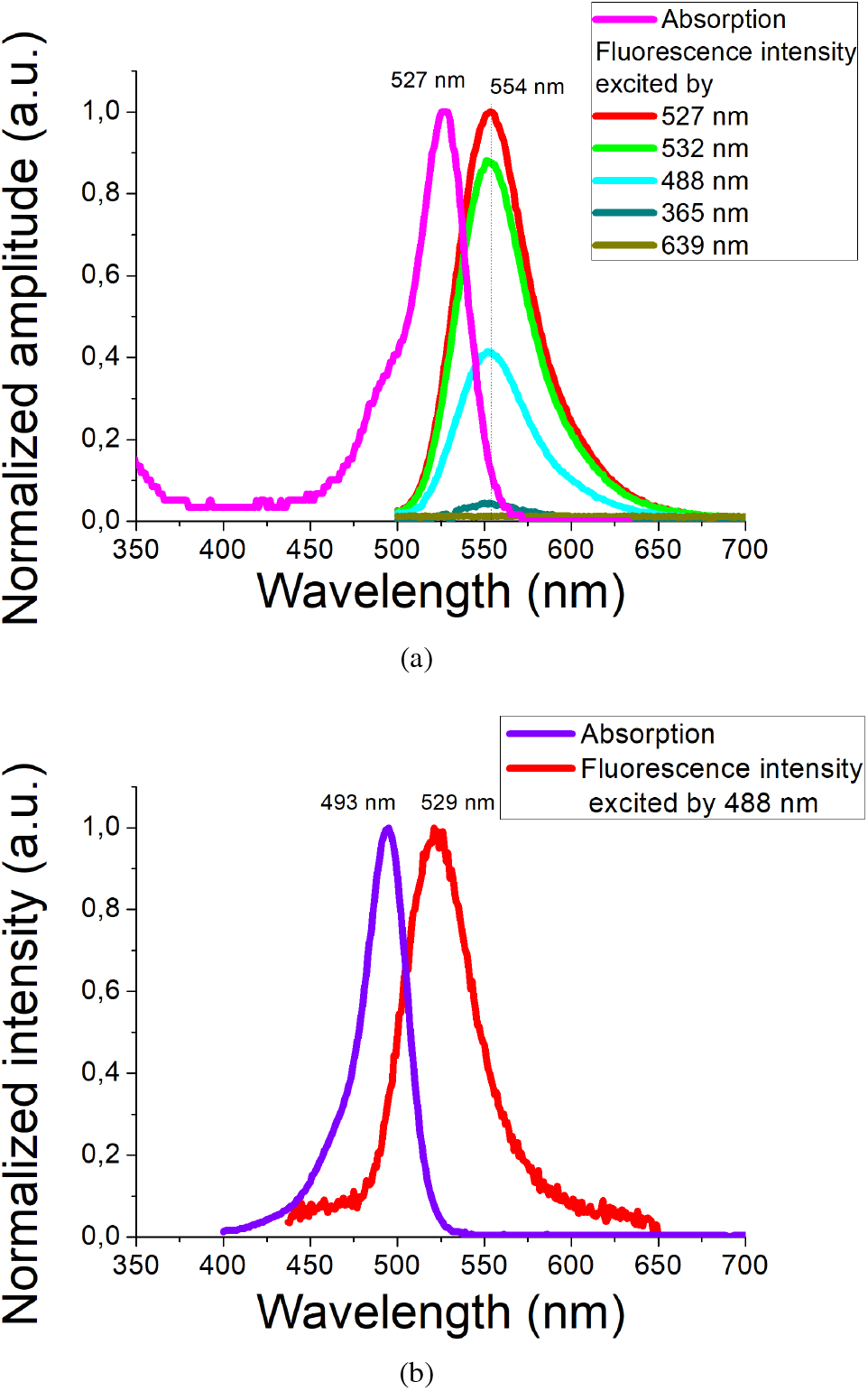
Rhodamine 6G (a) and DiBAC (b) emission and absorption spectra.

### 3.7. ABDS cost-effectivity

As mentioned earlier, one of the most important features of ABDS is its low-cost. Unlike expensive commercial cell dyes, ABDS can be manufactured in any laboratory and even in the field environment because the ingredients necessary for its manufacture – red permanent marker, ethanol, and PBS, are absolutely affordable. The cost-effectiveness of ABDS can be estimated using the following calculations. Edding pen 140S can be bought for $1.5 on Conrad.com or for $3 on eBay. In our laboratory we have checked that an approximately 200-m line can be drawn by using it, meaning that ABDS allows for 10^5^ staining cycles in $3 (see section 2.1). Contrary, 5 mg of DiBAC on MedChemExpress.com can be obtained for 70$. Since the molar mass of DiBAC is 516.63 g/mol [37], 5 mg of DiBAC approximately corresponds to 10 *μ*mol. DiBAC working solution has a concentration of 100 nM [49], so 5 mg of DiBAC can also be used for 10^5^ staining cycles, but their cost is 23 times higher than the cost of ABDS. Moreover, 100 *μ*L Deep Red Cell Mask can be purchased for $343 on Fishersci.com. This amount is sufficient for 100 cell staining cycles [50], which is significantly more expensive than ABDS. The price of Rhodamine 6G, which also stains cells very similarly to ABDS (Figure 5 and 6), varies greatly depending on its intended use and manufacturer. For example, 25 mg of Rhodamine 6G also from Fishersci.com can be purchased for $298. According to our absorbance studies, ABDS working solution should contain around 0.1 *μ*g of Rhodamine 6G in 1 mL. Therefore, 25 mg of $298 Rhodamine 6G is sufficient for 2.5 × 10^5^ cell staining cycles, or 10^5^ staining cycles for $120, which is still more expensive than ABDS. Other inexpensive options of Rhodamine 6G exist on www.medchemexpress.com (50 mg per $25), Carlroth.com (5 g per $30), and gtilaboratorysupplies.com (25 g per $30). At first glance, 25 g of Rhodamine 6G for $30 seems more efficient than 10 marker pens for $3, since the former allows to stain 250 times more probes. However, it should be noted that a permanent marker can last 91 years, even if three experiments with 1 ml of staining solution are performed daily. Thus, for small research groups, it is more cost-effective to invest in one marker than in 25 g of Rhodamine 6G, which they never use in full.

## 4. Conclusion

In this study, we developed a low-cost, open-source DIY cell imaging dye, called ABDS, based on permanent marker ink. We investigated the properties of ABDS, including emission, absorption, and Raman spectra (Figure 4), photobleaching, and biocompatibility (Figure 9). Our results indicate that ABDS is nontoxic and has properties similar to those of the fluorophore Rhodamine 6G, which is still under intensive investigation. However, a more remarkable result of our study is the ability of ABDS/R6G dyes to stain the cytoplasmic membrane as well as the endoplasmic reticulum, which can be clearly separated in fluorescence images using digital processing (Figures 5, 6, and 7). To our knowledge, the ability of R6G to stain the membrane of eukaryotic cells has not yet been reported.

ABDS is compatible with both 532 nm and 488 nm excitation sources, and a method for removing its fluorescence from one of these channels is discussed in Section 3.3 as a means of eliminating spectral overlap between ABDS and other fluorophores. The proposed cell staining method can also be considered as a biological approach for the isolation of small amounts of rhodamine 6G as an alternative to complex and dangerous high-voltage devices [47]. The data presented in Figure 3 and in Sec. 3.6 demonstrate that ABDS can be used to obtain detailed cell images, which allows to study cell adhesion, proliferation activity, and bioelectrical properties, *etc*.

Although only a few examples of the use of ABDS are presented in this paper, they show that ABDS is a promising candidate for the role of a cheap and accessible dye that can be easily used in the field environment and educational work because it does not require either careful sample preparation or special storage conditions. Our study also shows that labeling culture-ware with red permanent markers can lead to unexpected cell staining. We hope that the method of intravital cell staining with ABDS proposed in this work will significantly accelerate the development and testing of new cytological systems, including biosensors and bioelectronic devices.

## Supporting information

Membrane Proof

## 5. Acknowledgments

The authors express their gratitude to Metelkina E. M., Dubina Ph. M., Naschekina Yu.A., Blinova M. I., and Dubina M. V. for comprehensive assistance and support. This study was carried out with the support of the Ministry of Science and Higher Education (Project № FSRM-2024-0001).

## A. Supplementary data

In this section we provide additional information about the properties of commercially available dyes.

